# Non-targeted screening method for detecting temporal shifts in spectral patterns of Methicillin Resistance *Staphylococcus aureus* and post-hoc description of peak features

**DOI:** 10.1101/2024.11.27.625753

**Authors:** Kapil Nichani, Steffen Uhlig, Victor San Martin, Karina Hettwer, Kirstin Frost, Ulrike Steinacker, Heike Kaspar, Petra Gowik, Sabine Kemmlein

## Abstract

Non-targeted methods (NTMs) using matrix-assisted laser desorption/ionization time-of-flight mass spectrometry (MALDI-TOF MS) show promise in bacterial resistance detection, yet temporal variations in spectral features pose significant challenges. These proteomic patterns, which characterize bacterial phenotypes and pathological functions, may vary over time due to bacterial adaptation, virulence, or resistance mechanisms, resulting in large prediction uncertainties and potentially degrading NTM performance. We present a comprehensive screening method to detect temporal changes in MALDI-TOF spectral patterns, demonstrated using methicillin-resistant and -susceptible *Staphylococcus aureus* (MRSA/MSSA) isolates collected over several years. Our approach combines convolutional neural networks (CNN) with statistical methods, including significance testing, kernel density estimation, and receiver operating characteristics for dataset shift detection. We employ Gradient-weighted Class Activation Mapping (Gradcam) for post-hoc feature description, enabling biochemical characterization of temporal changes. This analysis reveals crucial insights into the dynamic relationship between spectral data patterns over time, addressing key challenges in developing robust NTMs for routine applications.

## 1. Introduction

Non-targeted methods (NTMs) using matrix-assisted laser desorption/ionization time of flight mass spectrometry (MALDI TOF MS) are a promising set of methods that can be used for bacterial species identification [1], sub-species or strain differentiation [2], metabolic profiling [3], among others. By measuring the mass-to-charge ratio of ions produced from bacterial samples, such methods provide proteomic fingerprints in form of spectra, which can be used to (a) query against a database of reference spectra or (b) used for supervised learning techniques; the latter of which we exploit here to detect changes in spectral patterns of antibiotic-resistant bacterial strains.

The use of machine learning methods with MALDI TOF spectra has been a “well-trodden path” for some time now [4,5]. These computational methods excel at learning intricate spectral patterns that reflect biochemical variations associated with antibiotic resistance, offering a promising NTMs for distinguishing Methicillin-resistant *Staphylococcus aureus* (MRSA) from its susceptible counterpart (MSSA) [5,6]. This enhanced analytical capability could significantly improve diagnostic precision, ultimately enabling more informed clinical decision-making.

Current literature on studying antimicrobial resistance (AMR) using MALDI TOF is very wide and deep, but with conflicting evidence [7]. While several groups have demonstrated promising results in MRSA/MSSA discrimination [6,8–10], other investigations highlight insufficiency and uncertainty in this approach as a rapid clinical diagnostic method [11–13]. These disparate outcomes likely stem from the inherent complexity of spectral data collection and analysis. Despite standardized sample preparation and analytical measurement methods (e.g. instrument configurations), spectra acquired from different (a) locations and (b) time points, frequently pose significant challenges for integration into a unified predictive model. Focusing on the latter, as one of the main practical challenges of NTMs is that the peak features of the spectra may change over time, due to various factors such as genetic mutations [14], environmental conditions [15], experimental protocols, or instrument settings [12]. These changes may affect the performance and reliability of NTM for bacterial identification and AMR detection, as well as the interpretation and comparison of the results across different studies and laboratories [6,16]. Most recently, this topic of data heterogeneity has been studied up Park and colleagues where they report effects of MRSA-MSSA class imbalance, effects of different sample preparation and laboratory environment for measurements [17]. This highlights the need to understand and account for the temporal changes in spectral peak features and understand how they impact the method performance.

Temporal changes in MALDI-TOF data manifest through multiple, interrelated mechanisms. Studies often employ potentially confusing terminology such as (a) concept drift, (b) covariate shift and (c) feature shift [18,19]. In practice, an example for (a) where new resistance patterns emerge or bacterial subgroups form; (b) when hospital-trained models face veterinary samples; and (c) where specific spectral patterns lose their discriminatory power. These terms, while theoretically distinct, often create confusion as their practical manifestations frequently overlap [20,21]. Rather than formally categorizing these phenomena, this work examines their collective impact through three interconnected aspects (see **Figure 1A**). First, the emergence of new resistance patterns could introduce novel spectral signatures, potentially compromising existing classification models. Second, shifts in bacterial population structure, where different resistance patterns becoming more prevalent over time, might affect model performance through changing class distributions. Third, specific peaks of interest (corresponding to specific protein complexes) may exhibit drift or instability, due to either instrumental factors or genuine biological changes in the bacterial proteome. Understanding these aspects is crucial for long-term implementation, as changes in any dimension could impact NTM reliability.

**Figure 1:**
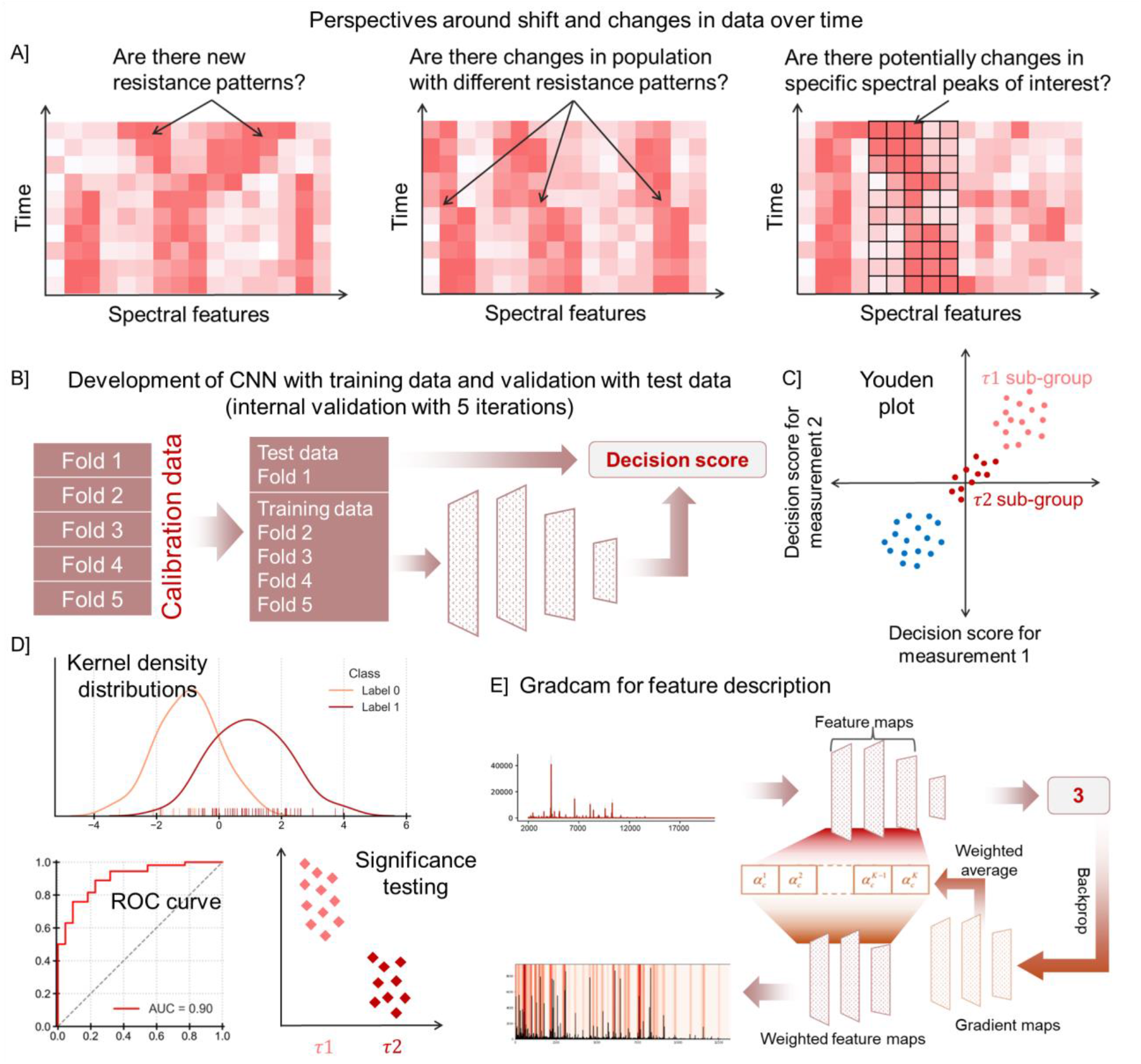
Schematic of the non-targeted screening method. (A) Different perspectives for data shift. (B) CNN training and testing, (C) Youden plot for decision scores, (D) Illustrative KDE, ROC curve and significance testing, (E) Gradcam for feature description.

Through analysis of MALDI-TOF spectral data from MRSA and MSSA isolates collected over several years, this paper addresses the challenge of temporal variation in spectral features. Our proposed method comprises three main steps, each targeting the key questions of resistance patterns, population changes, and spectral stability. First, we detect temporal changes using convolutional neural networks (CNN) trained on the complete dataset over entire time span (**Figure 1B)**, followed by a sequential approach to assess the significance of temporal differences in decision scores. Second, we conduct period-specific CNN training and classification to observe changes within MRSA. These shifts may not directly influence MRSA vs. MSSA classification but are essential for understanding intra-MRSA variations. By independently classifying different periods, we aim to further uncover changes in spectral patterns and target labels. The second step helps to detect the dataset shifts using the decision scores (**Figure 1C)**. Here we present the use of statistical procedures like visualizing kernel density estimation (KDE) distributions and receiver operating characteristics (ROC) curve (**Figure 1D)**. Lastly, we describe which set of peaks are contributing to temporal differences, using post-hoc feature description. Here, Gradient-weighted Class Activation Mapping (Gradcam)[22] is employed to identify relevant m/z ranges for each year (**Figure 1E)**. Extracting suitable peak features that contribute to the discrimination of a spectrum is not only important for the interpretability and performance of the models but also helps to detect biochemical changes. Post-hoc feature description helps to find distinctive patterns and features that lead to a classification decision. Ultimately, this comprehensive analysis enhances our understanding of the dynamic relationship between MRSA and MSSA across various years.

## 2. Materials and Methods

### 2.1 Spectral dataset

MALDI-TOF MS data were collected between 2008 and 202*1* at the Bundesamt fuer Verbraucherschutz und Lebensmittelsicherheit using a Bruker Microflex LT instrument. The dataset comprises spectra from 440 isolates of *Staphylococcus Aureus*. Sample preparation followed standard protocols [8,23]. Briefly, bacteria were cultured on blood agar plates for 24h at 37°C, transferred using a sterile loop to the MALDI target plate, overlaid with α-cyano-4-hydroxycinnamic acid matrix solution, and air-dried. Spectra were acquired in the mass range of 2,000-20,000 Da. Using custom scripts, the raw spectra underwent baseline extraction using rolling median.

### 2.2 Convolutional neural network (CNN) model training and decision scores

Two sets of CNN model training were performed – (i) classify MRSA and MSSA, (ii) classify time periods. For both, the CNN architecture comprised three convolutional layers interspersed with max-pooling layers, implemented via Keras and TensorFlow [24,25]. To address class imbalance across years, weighted training using Keras’s class weights function was applied, with weights inversely proportional to class frequencies, meaning that years and classes with fewer spectra have higher weights and vice versa. This way, the model gives more importance to the underrepresented years and classes and learns to generalize better across the data.

Nested cross-validation (NCV) approach utilized a calibration set consisting of MRSA and MSSA spectra in duplicates [26]. For each internal validation fold, the calibration set was divided into training and testing subsets. Adhering to the NCV framework, CNN models were trained on the training subset and validated with the test subset. CNN employed gradient descent optimization to minimize the loss function by iteratively adjusting the learnable parameters through backpropagation until convergence at each layer. The output from CNN comprised a standardized logit transformed probability value which was used for decision score computation [26]. The classification decision hinged on a threshold, set to zero in this context, such that positive decision scores indicated class A, while negative scores indicated class B.

### 2.3. Performance characteristics

Dual approaches were employed to evaluate the classifier’s performance. First, a confusion matrix quantified classification outcomes through true/false positives (TP/FP) and negatives (TN/FN). Second, the decision scores were analyzed to decompose variance components, providing deeper insights into classification reliability.

For binary non-targeted methods, the evaluation framework integrated both discrete classification metrics and continuous decision scores. Following Uhlig et al. [27,28], let *Z*_*ij*_ denote the decision score for isolate *i*, spectrum *j* [4]. The model for classification based on quantitative scores is as follows:

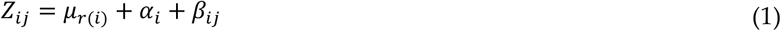

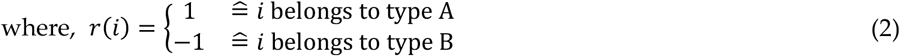

Here, *μr*(*i*) represents the mean score for respective classes A and B. *αi* captures the isolate *i* specific (random) variation and *β*_*ij*_ accounts for the (random) deviation of the decision score for spectrum *j* of isolate *i*. Assuming homogeneous variance within classes, two-way random effects analysis of variance ANOVA was used to estimate variance components.

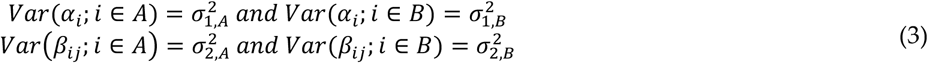

Based on the above variance components, the classification variances were calculated as follows:

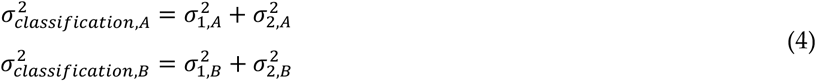

The smaller the classification variances, the better the classification power. The discrimination power *𝒟𝒫* derived from the concept of Fisher’s discrimination index of the NTM was then calculated as:

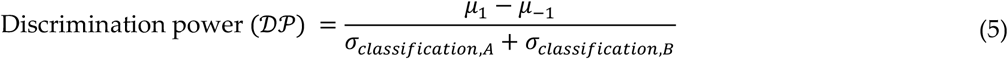

*σ*_*classification*_ in the denominator were replaced by estimated standard deviation values s_*classification*_ evaluated from the decision scores. Calculation of variance components was done according to ISO 5725-2 with PROLab statistical software (QuoData GmbH). *𝒟𝒫* above 2 means good discrimination between the classes. If *𝒟𝒫* is between 1 and 2, then reasonable discrimination is possible and if *𝒟𝒫* is less than 1, then poor discrimination.

### 2.4 Significance testing and t-values

The sequential t-test analysis was performed on the decision scores to assess significant differences between time periods (*τ*). At each sequential step, two time periods *τ*1 *and τ*2 were compared. If no significant differences were found, the periods were combined for the subsequent steps.

In case of unequal variance of decision scores, t-values for the two-time spans *τ*1 and *τ*2, were calculated as ratio of difference of the means 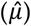 divided by pooled variance.

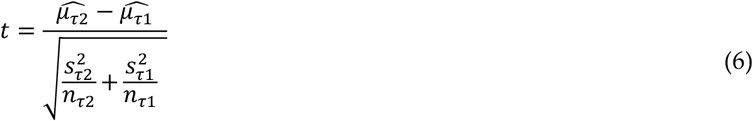

ROC curves were plotted and area under curve (AUC) were evaluated. A large AUC value suggests that the two time periods can be very well distinguished, with a value close to 1 signifying near perfect discrimination.

### 2.5 Feature extraction using Gradcam

Gradcam is a feature extraction technique based on neural network dissection, which aims to interpret the internal representations learned by deep networks [22]. It is based on the idea of using the gradients of the output class with respect to the feature maps of the last convolutional layer of the model, to produce a coarse localization map of the input [29]. Feature maps are the output of a convolutional layer in a neural network, which consists of a set of activation values for each filter (or kernel) applied to the input [30]. **Figure 1E** shows how Gradcam evaluation involved four key steps: First, prediction output was obtained from CNN in form of the decision score for the class of interest (say MRSA). Next, the gradient of the score *SR* with respect to the feature maps *A*_*convolution layer*. *l*_ of the last convolutional layer was evaluated. This was followed by performing a global average pooling on the gradients to obtain the weights 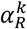 for each feature map. These weights represented the importance of each feature map for the MRSA. Each feature map *A*_*convolution layer*. *l*_ was then multiplied by its corresponding weight 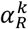 and summed up to get the Gradcam activation map *L*_*R*_. This map has the same dimension as that of the feature maps, *A*_*convolution layer*. *l*_. Lastly, the Gradcam activation map *L*_*R*_ is resized to match the size of the input image and overlay it on the original image with a color map. The resulting image shows the regions that CNN focuses on when predicting the class MRSA.

### 2.6 Peak features changing over time

After having identified the peak features, next to identify the specific peak features that change over the years, we perform a one-way ANOVA analysis. We calculated the *F*-value for the features, which can be used to determine the features that contribute most to the classification. By performing an ANOVA analysis on the Gradcam feature values, we can test whether there are significant differences among the mean values of each peak for each year. These peaks are the ones that have changed significantly over the years and can be considered potential biomarkers or indicators of temporal variation.

## 3. Results

### 3.1 Identifying temporal changes using classification decision scores for MRSA and MSSA

To address the key objective of identifying spectral shifts, first CNNs were trained using all available spectral data for MRSA and MSSA, i.e. from 2008 to 2021 employing the 5-fold NCV technique. Samples from each year were evenly distributed across all folds to ensure an equal representation of years. Internal validation decision scores were evaluated and no spectra for external validation were used as the aim was not to develop a NTM to predict MRSA or MSSA, rather to gain insights into the varying spectral patterns of MRSA and MSSA, that affects their classification. In doing so one can detect if there are changes in the population with different spectral patterns.

**Figure 2** shows a Youden plot with decision scores for distinguishing MRSA and MSSA. Each point on the plot represents an isolate with decision scores for duplicate measurements on the two axes. Considering a decision threshold of 0, a confusion matrix can be constructed, resulting in 14 false positives and 14 false negatives, resulting in a true positive rate (sensitivity) of 91% and a false positive rate (1 — specificity) of about 3.3%. From the Youden plot, two direct observations emerge. First, for MSSA samples (spanning from 2008 to 2017) the classification remains predominantly accurate, i.e., majority of the points lie in the third quadrant. And secondly, the decision scores of replicate samples display considerable variability, as evidenced by their deviation from the line of identity.

**Figure 2:**
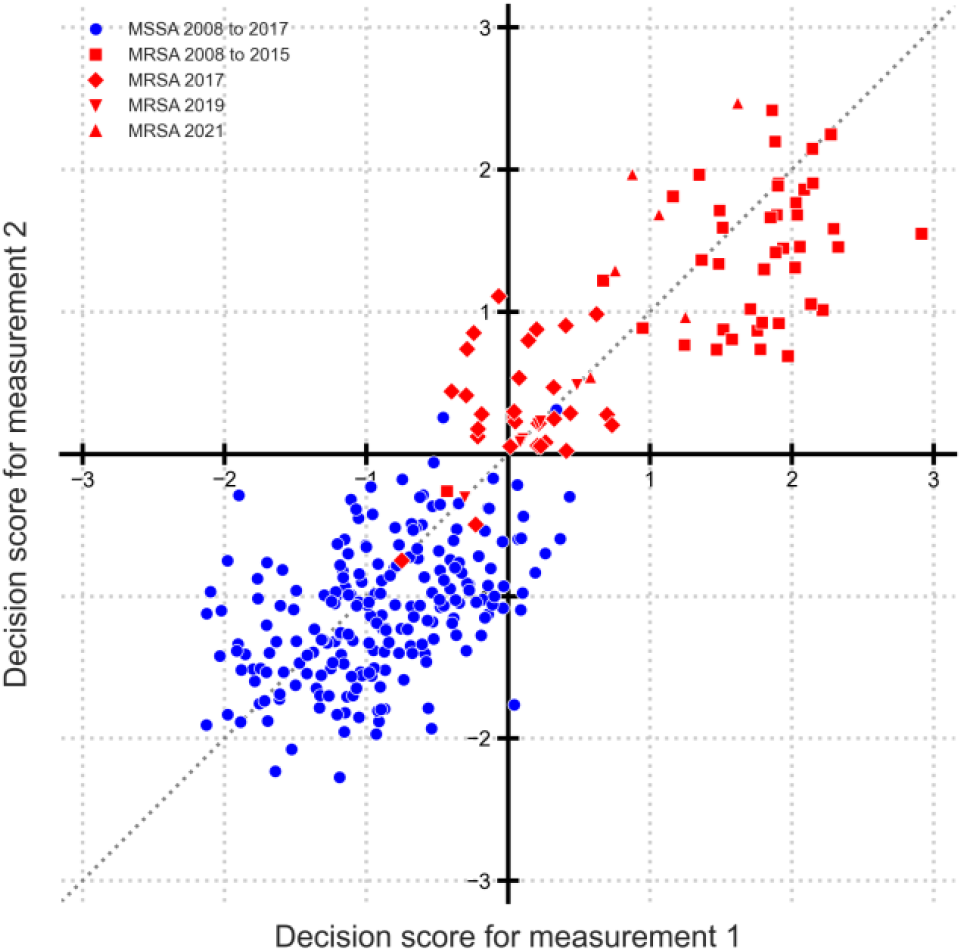
Decision scores of MRSA vs MSSA as Youden plot. Blue circles show MSSA from 2008 to 2017. Red squares show MRSA from 2008 to 2015. Red diamonds show MRSA from 2017. Red lower triangles show MRSA from 2019 and red upper triangle from 2021. Line of identity is shown as dotted line.

To examine the presence of “sub-populations” within MRSA, the point clouds are drawn in different shapes by known year label. MRSA samples from 2008 to 2015 (red squares) and 2021 (red upper triangle), are farthest from the MSSA of 2008 to 2017 (blue circles), which implies that they can be more distinctly classified. The point cloud distribution additionally indicates that MRSA samples from 2017 (red diamonds) and 2019 (red lower triangle) are notably close to the MSSA samples. This suggests that clear classification between MRSA samples from these years is challenging. There is also indication that 2019 is similar to 2017, but 2021 is different. However, further differentiation is not possible, i.e., comparing 2021 with other years, because of the fewer available data points.

Such a result prompts the question of how one can identify evolving patterns or spot temporal variations within the data over time. In order to identify whether differences in subgroups are significant, we perform a sequential ttest procedure.

### 3.2 Sequential significance testing of decision scores

Following the observations from the Youden plot, sequential significance testing to systematically identify temporal groupings was performed. **Figure 3** illustrates the step-by-step approach. The sequential testing began with step 1, comparing two pairs of years with minimal data points: 2019 versus 2021 (denoted by red 1) revealed statistically significant differences, while 2008 versus 2009 (marked as green 2) showed no significant differences. Following this initial analysis, 2008 and 2009 were combined for subsequent comparisons. Throughout the figure, green-shaded boxes indicate pairs of periods where decision scores show no statistical differences, while red-shaded boxes highlight periods with statistically significant differences.

**Figure 3:**
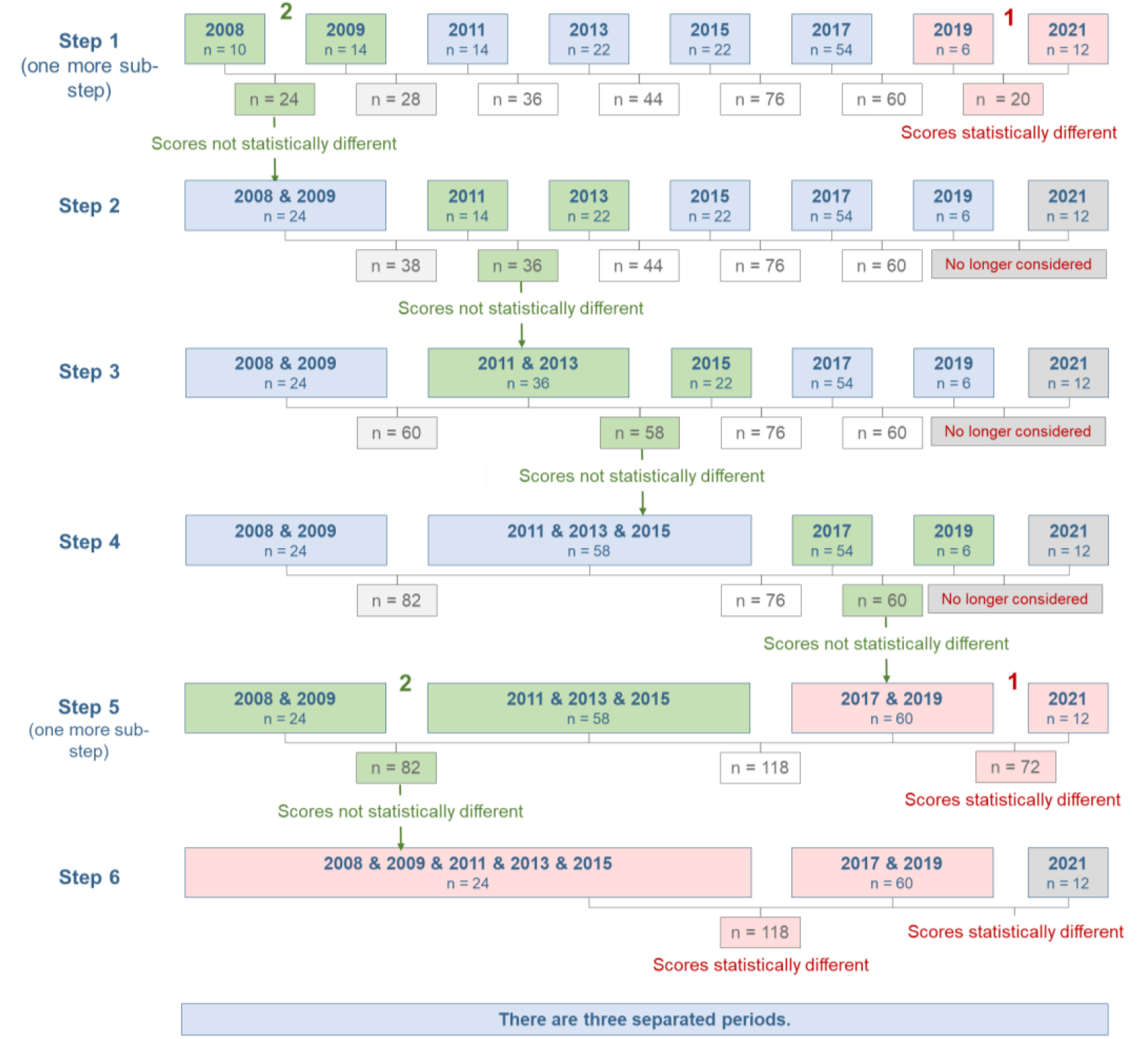
Sequential stepwise testing for differences in scores for MRSA. This schematic illustrates the iterative process of comparing and merging time periods based on their decision scores. In step one, pairs of periods with the smallest sample sizes were compared. Green-shaded boxes indicate periods with statistically similar scores that were merged for subsequent analysis, while red-shaded boxes show periods with statistically different scores that remained separate. When two comparisons occurred within the same step, they are labeled as ’1’ and ’2’. Through this sequential comparison and merging of statistically similar periods, three distinct time periods emerged with significantly different decision scores.

Step 2 revealed that decision scores from 2011 and 2013 showed no significant differences. Proceeding to step 3, the comparison of the combined 2011+2013 period with 2015 also showed no statistical differences, enabling the formation of larger temporal groups. Further analysis demonstrated that decision scores from 2017 and 2019 were statistically similar, while the data from 2008 to 2015 maintained statistical consistency throughout the analysis.

The sequential testing culminated in step 6, which definitively identified three distinct temporal periods: 2008-2015, 2017-2019, and 2021. Final statistical tests confirmed significant differences between these three periods. The sequential analysis provides strong evidence that MRSA isolates from 2017 and 2019 exhibit similar characteristics but differ significantly from other periods. While this methodology effectively addresses the presence of changes in spectral patterns, it is important to note that these statistical differences cannot be directly interpreted as qualitative differences in MRSA isolate behavior.

### 3.2 Finding breaks by year wise-training

The sequential significance testing helped identify three time periods from the decision scores. To further validate this result, several CNNs were trained to perform classification for two time periods, separately for MRSA and MSSA. **Figure 4** shows the (A) rug plot charts for decision scores, showing the distribution of scores and (B) ROC curves showing discriminatory relation of the model through the lens of false-positive rate (FPR) and true positive rate (TPR). Calculating the t-values using Equation 6, we obtain large t-values when comparing MRSA for periods (a) 2015 versus 2017 (value of 6.9), and (b) 2017 versus 2019+2021 (value of 4.67). Considering a critical t-value of 2, one can consider the difference in the decision scores to be significant for the above two periods. The ROC curves also show high discrimination for these two periods with AUC values of 0.9 and 0.83, respectively. For the other periods, the AUC value is around 0.5.

**Figure 4:**
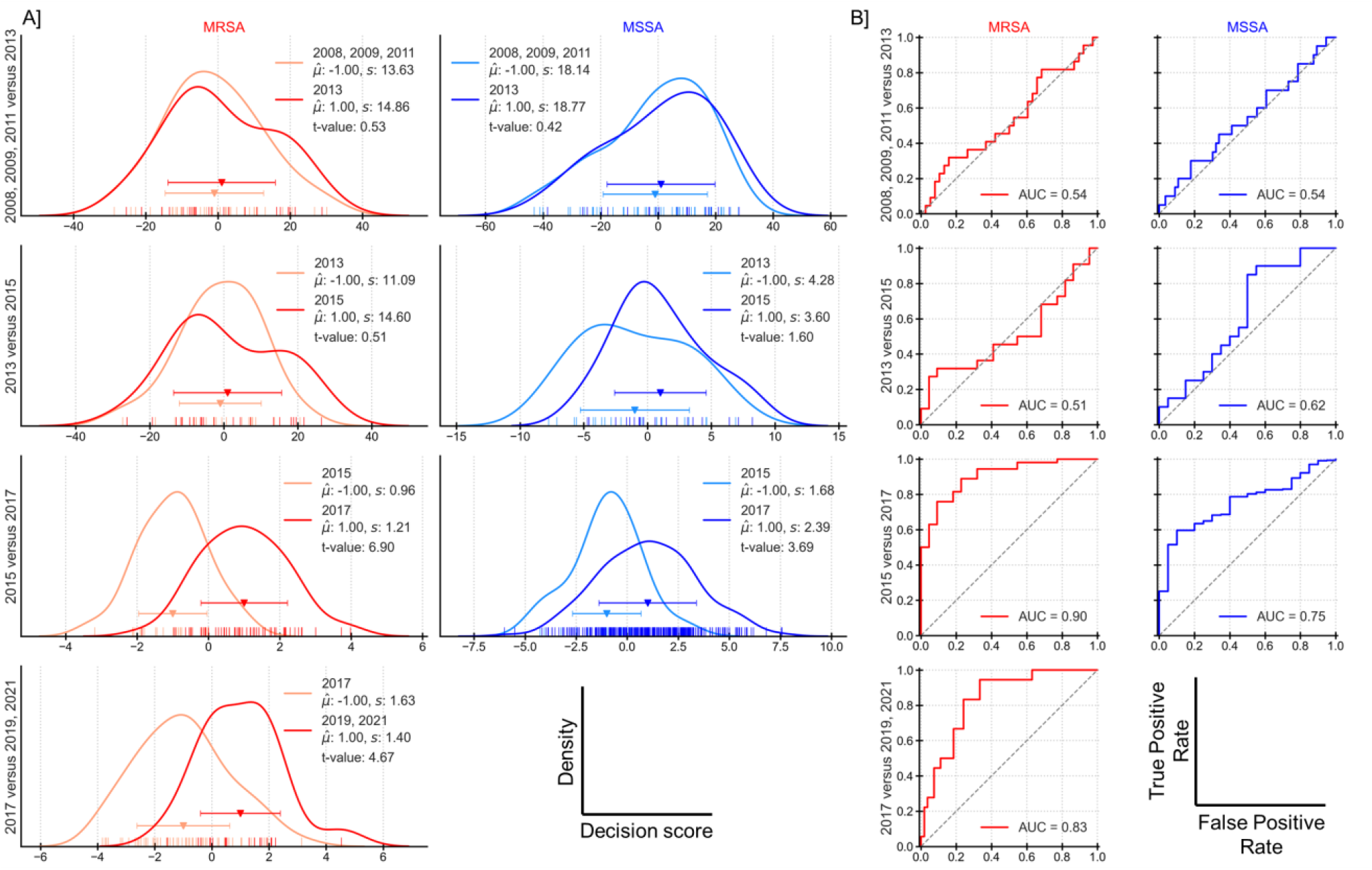
Distinction between the different time periods separately for MRSA and MSSA depicted in rows. (A) Rug plots with kernel density estimation curves. Plotted are the mean and standard deviation. (B) ROC curves with AUC values for the classification.

Likewise, MSSA for period 2015 versus 2017 also shows some evidence of differences in decision scores with relatively high t-value of 3.69 and AUC of 0.75. This finding is particularly intriguing as antibiotic resistance-associated proteins should theoretically not influence MSSA spectra. Several potential explanations warrant further investigation: the presence of borderline resistance cases that are still classified as MSSA, regional and temporal variations in MSSA subtypes across different geographical regions, differences in measurement conditions during spectral acquisition, or variations in isolate processing protocols that could affect bacterial viability and proteome expression. The combination of t-values and ROC-AUC metrics provides complementary evidence: t-values establish statistical significance of temporal differences, while ROC-AUC values quantify classification performance. These metrics jointly support the temporal breaks identified in the previous section.

### 3.3 Tracking features over time

#### 3.3.1 Feature extraction heatmaps

We proceeded to neural network dissection with Gradcam to address the question if there are changes in specific peaks. The left panels of **Figure 5A** show the baseline-corrected spectra (black line) overlaid the Gradcam heatmaps presenting one representative spectrum from each year. These heatmaps quantitatively illustrate the contribution of each spectral region to the model’s predictions, where warmer colors indicate stronger contributory features. For instance, in the spectrum in the first row, multiple specific bands demonstrate high influence on the classification. Notably, across all spectra, regions corresponding to m/z 2000-4000 consistently show high importance, indicated by prominent dark red bands. Several studies also report presence of discriminatory peaks in that range [5,7,31].

**Figure 5:**
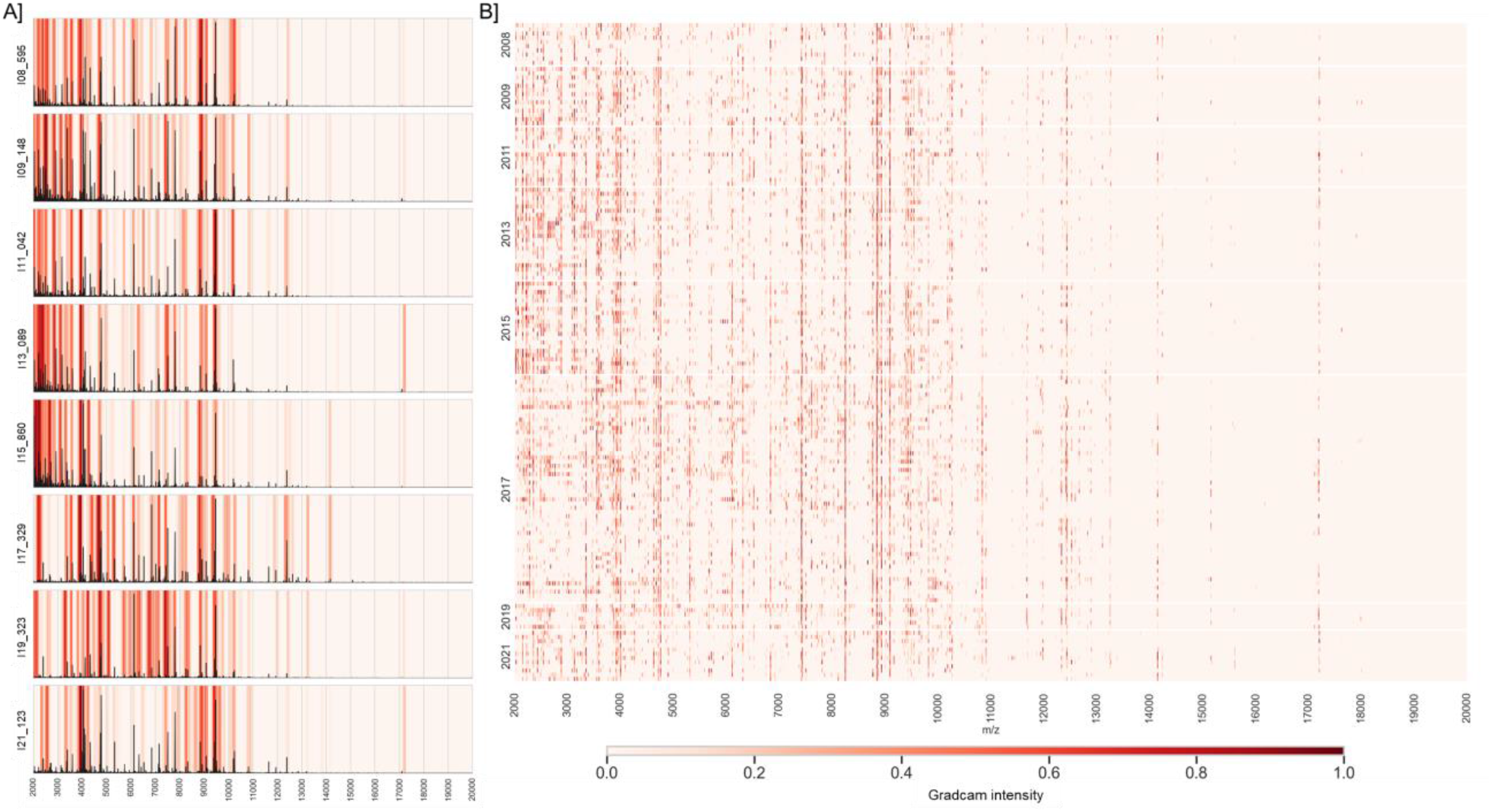
Gradcam feature values (A) exemplary for selected spectra and (B) plotted as a heatmap for all spectra as a heatmap.

By bringing together the Gradcam results for all the spectra, a consolidated heatmap is constructed in **Figure 5B**. In this heatmap, each row of pixels corresponds to a spectrum and each column corresponds to a mass by charge ratio. Once again, the color intensity represents the classification importance derived from Gradcam analysis, with warmer colors indicating higher significance. The heatmap reveals distinct patterns of CNN focus across different years, suggesting temporal variations in MRSA spectra.

Overall, Gradcam demonstrates its utility as a sophisticated visualization technique for interpreting CNN classification behavior, offering insights into the network’s learned features and their relationship to output classes across varying inputs. It can also be used for development and optimization of NTMs. The heatmap primarily provides a qualitative or global perspective of the network-learned spectral features, and detailed year-to-year comparisons may only be partially feasible through visual inspection.

#### 3.3.2 Detecting specific peaks sensitive to temporal changes

To quantitatively evaluate the year-to-year changes in the Gradcam features, one-way ANOVA analysis is performed for the Gradcam feature value with the year. High F-values mean that the peak regions show large variation in importance between the years year. Six exemplary spectral regions are taken with relatively high F-value and plotted in **Figure 6**. The overlap of features learned by the models with recognized resistance patterns reported in literature is a promising result. Further work is warranted here in terms of biochemical characterization of the proteins responsible for the peaks.

**Figure 6:**
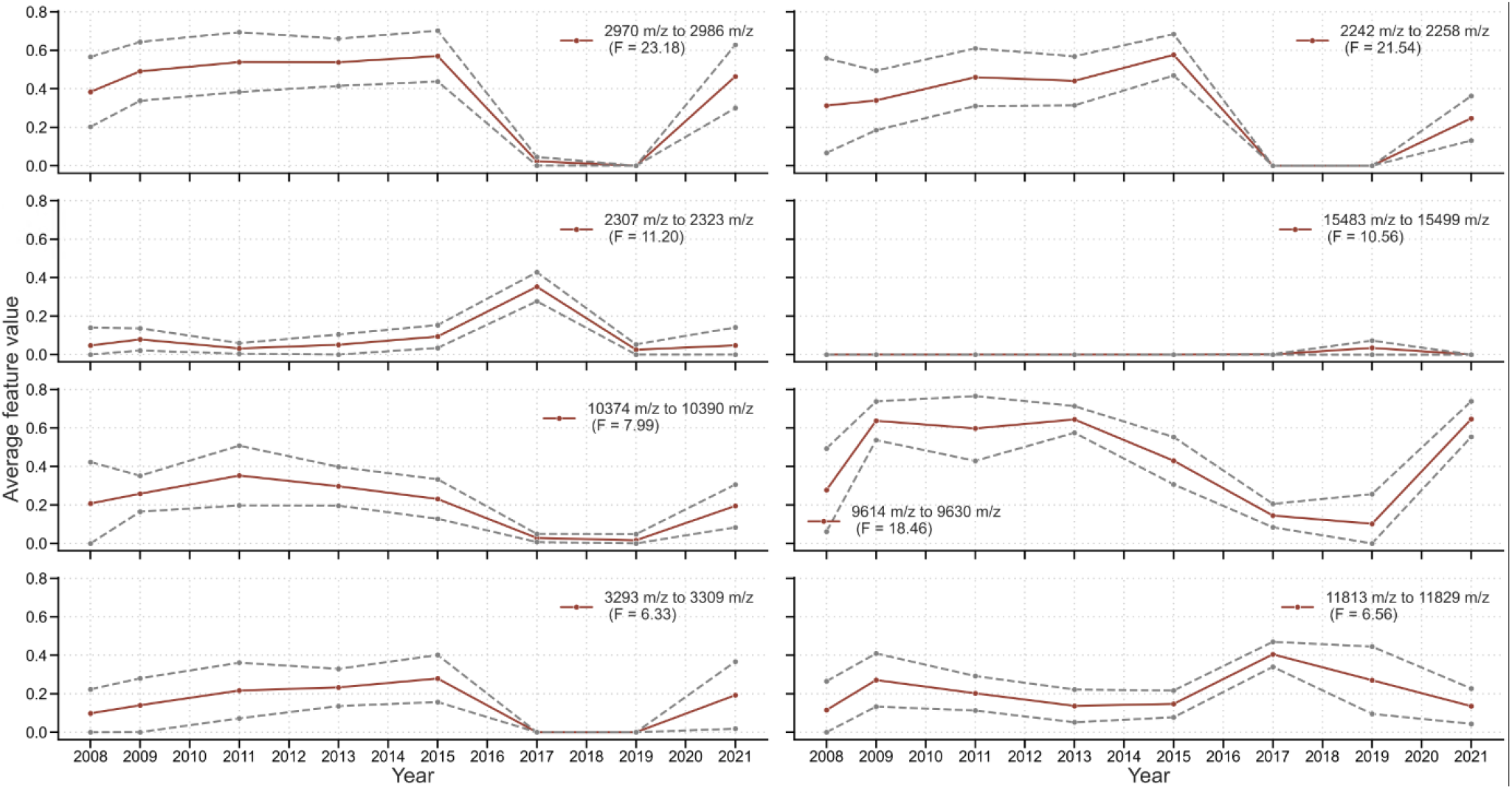
Exemplary patterns of Gradcam feature values changing over time for size m/z ranges.

### 3.4 Classification performance based on different time periods

Following the identification of temporal breaks in 2017, we investigated classification performance by analyzing pre-2017 and post-2017 periods, separately. **Figure 7** presents Youden plots for MRSA classification across these distinct periods: (A) 2008-2015 and (B) 2017-2019. The 2008-2015 period demonstrated a TPR, or sensitivity of 99% and very low FPR (<<1%), i.e. very high specificity. This represents a substantial improvement over the non-temporally segregated analysis presented in Section 3.1. The repeatability standard deviations for MRSA and MSSA are 0.404 and 0.285, respectively, with classification standard deviations of 0.419 and 0.308, respectively. Using Equation 5, *𝒟𝒫* is calculated to be 2.75. Recollect, that values above 2 means very good discrimination between the classes. The 2017-2019 period showed markedly different characteristics, with sensitivity decreasing to 52% (FPR 11%). Further, it is also clear that the variation between the replicates has increased (spread of the red and blue points), with the repeatability standard deviations for MRSA and MSSA of 0.815 and 0.729, and classification standard deviations of 0.815 and 0.821. The resulting *𝒟𝒫* of 1.22 indicates that the performance characteristics for the NTM for identifying MRSA from MSSA has significantly deteriorated, necessitating a revision and update of the NTM [32].

**Figure 7:**
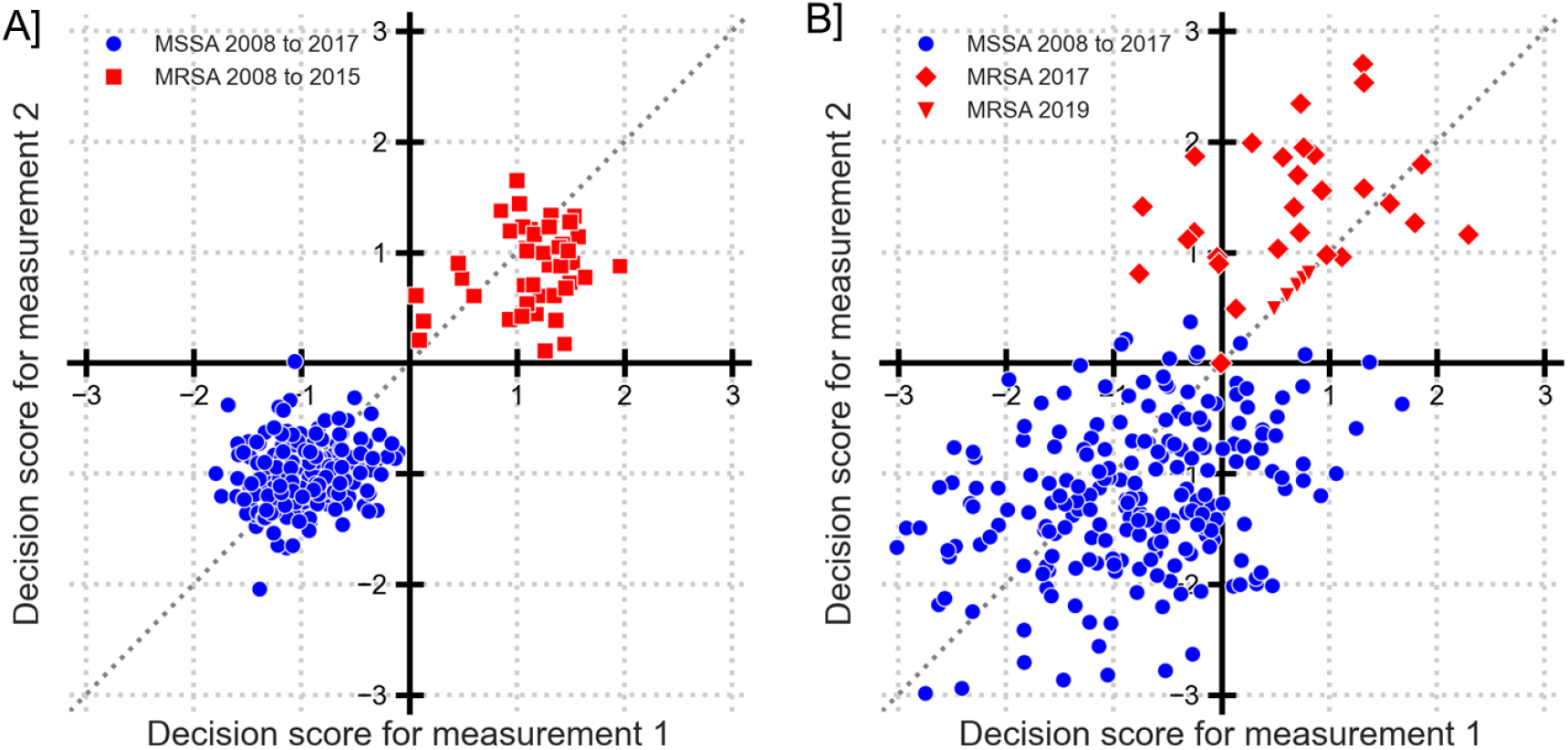
Youden plot for discrimination of MRSA and MSSA with the split time periods. (A) MSSA from 2008 to 2017 (blue dots) and MRSA from 2008 to 2015 (red square). (B) MSSA from 2008 to 2017 (blue dots) and MRSA from 2017 (red diamonds) and from 2019 (red lower triangles).

## 4. Discussion

Proteomic patterns, such as protein expression, modification, and interaction, are widely used to characterize the biological and pathological functions of bacteria. However, these patterns may vary over time due to various processes, such as bacterial growth, adaptation, virulence, or resistance. In this paper, we present a study of the temporal changes in proteomic patterns of MRSA and propose a screening method. We use a large sample of MALDI TOF data collected over several years. We propose and apply statistical methods and machine learning techniques to find suitable features and track their changes over the years. We also discuss the implications of our results and how the identified peaks can help with biochemical characterization as well as the prospects for future studies with emerging technologies.

We use CNN to learn cross-year feature representations that can capture the biochemical variations associated with antibiotic resistance or susceptibility. We evaluate our method on a large dataset of mass spectrometry data collected from 2008 to 2021. Quantitative decision scores from CNNs were evaluated and used for not only performance characterization, but also to evaluate the changes over the years. Using decision scores, variance components are evaluated from which the precision parameters are calculated such as repeatability standard deviation and classification standard deviation, can help to describe the distribution of points and the farness to each other. Detecting shifts in data over years using quantitative decision scores is an important novelty of this work, which would not have been possible merely with qualitative binary outcomes.

Feature descriptions are primarily important for 3 main reasons. First, to ensure that the method works “as expected”, feature extraction can help understand if the model identifies features that are “logical” or “relevant”, or if the model relies on arbitrary patterns in the data to make a classification. Second, to understand the reason for the result, one tries to understand why the model classified the sample as MRSA. And more importantly, how can we determine whether the features have changed over the years. Lastly, to develop targeted methods or improved NTMs. Neural network dissection methods like Gradcam are a type of post-hoc feature description method that analyze the activations of the hidden layers of a network and try to find meaningful associations between the units in those layers and the input features or the output classes. We apply the Gradcam method to detect the relevant mass ranges for each year and compare them with the temporal breaks detected by statistical methods. We show that our approach can provide useful insights for biochemical analysis, as well as potentially accounting for the possible confounding factors of sample selection, isolate preparation, and measurement circumstances.

Traditional accuracy-based performance metrics (where accuracy = (TP+TN) / (TP+FP+TN+FN)) might be misleading when dealing with extremely imbalanced data, where one class considerably dominates the other(s). They are commonly reported in literature where new NTMs are described. The model in the NTM can simply predict the majority class and have a large accuracy. Here there is a risk of having a false sense of superior NTM performance.

It is advisable to exercise caution or, ideally, refrain from employing accuracy metrics commonly applied for model evaluation when faced with imbalanced class structures, as their interpretation may yield misleading insights. Quantitative decision scores should rather be used, as done here from which sensitivity and specificity can be evaluated. The work herein limits by all means to a screening method and does not offer any confirmatory procedures because so far it does not provide any information on the biochemical nature of the changed patterns or extracted features. Nevertheless, the approaches offer a pragmatic and powerful approach that can be implemented via use of simple tools as shown in this work. Therefore, further biochemical analysis is needed to identify and describe the proteins responsible for the peaks. The core reasons for the changes need further investigation. Here, another possible reason is the different sampling process of the isolates, which can also affect how the spectra look. Overall, it is clear that with such a screening method, one can easily detect changes in data and generate several hypotheses to further confirm.

## 5. Conclusions

Data shifts and change in the samples over time is a huge challenge for any NTM for species typing and AMR programs, let alone other analytical sciences. And this work helps to provide tools for handling such scenarios. In this paper, we propose a non-targeted screening method to detect the changes of peak features in mass spectrometry data over the years. We have shown that by identifying the temporal breaks, strategic dataset selection and model adaptation, one can maintain classification accuracy across different time periods.

Performance evaluation of a non-targeted method is very important not only to trust the results, but also for gaining wider acceptance in routine use. We show how the performance evaluation can be done using the underlying decision scores obtained from the CNN models. In this work, several data experiments were conducted with MRSA and MSSA MALDI TOF spectra. To investigate the impact of changing samples, different combinations of available data were put together to test out hypotheses. The endeavor of NTM development encounters heightened challenges when the MRSA (or any class of interest) contends with a limited pool of unique available samples, compounded by evolutionary changes in features across temporal dimensions. This is something we expect in reality and hence the work in this report describes considerations and options to tackle these challenges. Lastly, the approaches described, when provided in a suitable environment add to the toolbox for developing superior methods of measurement and validating them. Future work should focus on implementing the screening method described as an automated method for detecting spectral drift and implementing real-time model updates to ensure reliable clinical diagnostics.

## References

1. Wolters, M.; Rohde, H.; Maier, T.; Belmar-Campos, C.; Franke, G.; Scherpe, S.; Aepfelbacher, M.; Christner, M. MALDI-TOF MS Fingerprinting Allows for Discrimination of Major Methicillin-Resistant Staphylococcus Aureus Lineages. International Journal of Medical Microbiology 2011, 301, 64–68, doi:10.1016/j.ijmm.2010.06.002.

2. Pérez-Sancho, M.; Vela, A.I.; Horcajo, P.; Ugarte-Ruiz, M.; Domínguez, L.; Fernández-Garayzábal, J.F.; De La Fuente, R. Rapid Differentiation of Staphylococcus Aureus Subspecies Based on MALDI-TOF MS Profiles. J VET Diagn Invest 2018, 30, 813–820, doi:10.1177/1040638718805537.

3. Złoch, M.; Pomastowski, P.; Maślak, E.; Monedeiro, F.; Buszewski, B. Study on Molecular Profiles of Staphylococcus Aureus Strains: Spectrometric Approach. Molecules 2020, 25, 4894, doi:10.3390/molecules25214894.

4. Li, L.; Garden, R.W.; Sweedler, J.V. Single-Cell MALDI: A New Tool for Direct Peptide Profiling. Trends in Biotechnology 2000, 18, 151–160, doi:10.1016/S0167-7799(00)01427-X.

5. Tang, R.; Luo, R.; Tang, S.; Song, H.; Chen, X. Machine Learning in Predicting Antimicrobial Resistance: A Systematic Review and Meta-Analysis. International Journal of Antimicrobial Agents 2022, 60, 106684, doi:10.1016/j.ijantimicag.2022.106684.

6. Weis, C.V.; Jutzeler, C.R.; Borgwardt, K. Machine Learning for Microbial Identification and Antimicrobial Susceptibility Testing on MALDI-TOF Mass Spectra: A Systematic Review. Clinical Microbiology and Infection 2020, 26, 1310–1317, doi:10.1016/j.cmi.2020.03.014.

7. Sauget, M.; Valot, B.; Bertrand, X.; Hocquet, D. Can MALDI-TOF Mass Spectrometry Reasonably Type Bacteria? Trends in Microbiology 2017, 25, 447–455, doi:10.1016/j.tim.2016.12.006.

8. Østergaard, C.; Hansen, S.G.K.; Møller, J.K. Rapid First-Line Discrimination of Methicillin Resistant Staphylococcus Aureus Strains Using MALDI-TOF MS. International Journal of Medical Microbiology 2015, 305, 838–847, doi:10.1016/j.ijmm.2015.08.002.

9. Yu, J.; Tien, N.; Liu, Y.-C.; Cho, D.-Y.; Chen, J.-W.; Tsai, Y.-T.; Huang, Y.-C.; Chao, H.-J.; Chen, C.-J. Rapid Identification of Methicillin-Resistant Staphylococcus Aureus Using MALDI-TOF MS and Machine Learning from over 20,000 Clinical Isolates. Microbiol Spectr 2022, 10, e00483-22, doi:10.1128/spectrum.00483-22.

10. Kim; Kim; Chung; Chung; Han; Kim Rapid Discrimination of Methicillin-Resistant Staphylococcus Aureus by MALDI-TOF MS. Pathogens 2019, 8, 214, doi:10.3390/pathogens8040214.

11. Paskova, V.; Chudejova, K.; Sramkova, A.; Kraftova, L.; Jakubu, V.; Petinaki, E.A.; Zemlickova, H.; Neradova, K.; Papagiannitsis, C.C.; Hrabak, J. Insufficient Repeatability and Reproducibility of MALDI-TOF MS-Based Identification of MRSA. Folia Microbiol 2020, 65, 895–900, doi:10.1007/s12223-020-00799-0.

12. Lasch, P.; Fleige, C.; Stämmler, M.; Layer, F.; Nübel, U.; Witte, W.; Werner, G. Insufficient Discriminatory Power of MALDI-TOF Mass Spectrometry for Typing of Enterococcus Faecium and Staphylococcus Aureus Isolates. Journal of microbiological methods 2014, 100, 58–69.

13. Lasch, P.; Wahab, T.; Weil, S.; Pályi, B.; Tomaso, H.; Zange, S.; Kiland Granerud, B.; Drevinek, M.; Kokotovic, B.; Wittwer, M.; et al. Identification of Highly Pathogenic Microorganisms by Matrix-Assisted Laser Desorption Ionization–Time of Flight Mass Spectrometry: Results of an Interlaboratory Ring Trial. J Clin Microbiol 2015, 53, 2632–2640, doi:10.1128/JCM.00813-15.

14. Vestergaard, M.; Frees, D.; Ingmer, H. Antibiotic Resistance and the MRSA Problem. Microbiology spectrum 2019, 7, 10–1128.

15. Viboud, G.; Asaro, H.; Huang, M.B. Use of Matrix-Assisted Laser Desorption Ionization Time of Flight (MALDI-TOF) to Detect Antibiotic Resistance in Bacteria: A Scoping Review. American Journal of Clinical Pathology 2024, 161, 317–328, doi:10.1093/ajcp/aqad160.

16. Kannan, E.P.; Gopal, J.; Muthu, M. Analytical Techniques for Assessing Antimicrobial Resistance: Conventional Solutions, Contemporary Problems and Futuristic Outlooks. TrAC Trends in Analytical Chemistry 2024, 178, 117843, doi:10.1016/j.trac.2024.117843.

17. Park, Y.; Weig, M.; Noll, C.; Bader, O.; Hauschild, A.-C. Effect of Data Heterogeneity in Clinical MALDI-TOF Mass Spectra Profiles on Direct Antimicrobial Resistance Prediction through Machine Learning 2024.

18. Kull, M.; Flach, P. Patterns of Dataset Shift. 2014.

19. Zhang, H.; Singh, H.; Ghassemi, M.; Joshi, S. “Why Did the Model Fail?”: Attributing Model Performance Changes to Distribution Shifts 2023.

20. Moreno-Torres, J.G.; Raeder, T.; Alaiz-Rodríguez, R.; Chawla, N.V.; Herrera, F. A Unifying View on Dataset Shift in Classification. Pattern Recognition 2012, 45, 521–530, doi:10.1016/j.patcog.2011.06.019.

21. Gama, J.; Žliobaitė, I.; Bifet, A.; Pechenizkiy, M.; Bouchachia, A. A Survey on Concept Drift Adaptation. ACM Comput. Surv. 2014, 46, 1–37, doi:10.1145/2523813.

22. Selvaraju, R.R.; Cogswell, M.; Das, A.; Vedantam, R.; Parikh, D.; Batra, D. Grad-Cam: Visual Explanations from Deep Networks via Gradient-Based Localization. In Proceedings of the Proceedings of the IEEE international conference on computer vision; 2017; pp. 618–626.

23. Werner, G.; Fleige, C.; Feßler, A.T.; Timke, M.; Kostrzewa, M.; Zischka, M.; Peters, T.; Kaspar, H.; Schwarz, S. Improved Identification Including MALDI-TOF Mass Spectrometry Analysis of Group D Streptococci from Bovine Mastitis and Subsequent Molecular Characterization of Corresponding Enterococcus Faecalis and Enterococcus Faecium Isolates. Veterinary Microbiology 2012, 160, 162–169, doi:10.1016/j.vetmic.2012.05.019.

24. Chollet, F.; others Keras 2015.

25. Abadi, M.; Barham, P.; Chen, J.; Chen, Z.; Davis, A.; Dean, J.; Devin, M.; Ghemawat, S.; Irving, G.; Isard, M.; et al. TensorFlow: A System for Large-Scale Machine Learning. 21.

26. Nichani, K.; Uhlig, S.; Colson, B.; Hettwer, K.; Simon, K.; Bönick, J.; Uhlig, C.; Kemmlein, S.; Stoyke, M.; Gowik, P.; et al. Development of Non-Targeted Mass Spectrometry Method for Distinguishing Spelt and Wheat. Foods 2022, 12, 141.

27. Uhlig, S.; Nichani, K.; Stoyke, M.; Gowik, P. Validation of Binary Non-Targeted Methods: Mathematical Framework and Experimental Designs. bioRxiv 2021, doi:10.1101/2021.01.19.427235.

28. Uhlig, S.; Nichani, K.; Colson, B.; Hettwer, K.; Simon, K.; Uhlig, C.; Stoyke, M.; Steinacker, U.; Becker, R.; Gowik, P. Performance Characteristics and Criteria for Non-Targeted Methods. In Proceedings of the Eurachem workshop; Tartu, Estonia, 2019.

29. Draelos, R.L.; Carin, L. Use HiResCAM Instead of Grad-CAM for Faithful Explanations of Convolutional Neural Networks. arXiv preprint arXiv:2011.08891 2020.

30. Zeiler, M.D.; Fergus, R. Visualizing and Understanding Convolutional Networks. In Proceedings of the Computer Vision– ECCV 2014: 13th European Conference, Zurich, Switzerland, September 6-12, 2014, Proceedings, Part I 13; Springer, 2014; pp. 818–833.

31. Shah, H.N.; Rajakaruna, L.; Ball, G.; Misra, R.; Al-Shahib, A.; Fang, M.; Gharbia, S.E. Tracing the Transition of Methicillin Resistance in Sub-Populations of Staphylococcus Aureus, Using SELDI-TOF Mass Spectrometry and Artificial Neural Network Analysis. Systematic and applied microbiology 2011, 34, 81–86.

32. Nichani, K.; Uhlig, S.; Stoyke, M.; Kemmlein, S.; Ulberth, F.; Haase, I.; Döring, M.; Walch, S.G.; Gowik, P. Essential Terminology and Considerations for Validation of Non-Targeted Methods 2022.

